# Development of novel potent ligands for GPR85, an orphan G protein-coupled receptor expressed in the brain

**DOI:** 10.1101/2021.09.18.460934

**Authors:** Aya Sakai, Takeshi Yasui, Masashi Watanave, Rine Tatsumi, Yoshihiko Yamamoto, Wataru Takano, Yuki Tani, Izumi Okamura, Hirokazu Hirai, Shigeki Takeda

## Abstract

GPR85 is a member of the G protein-coupled receptor and is a super-conserved receptor expressed in the brain sub-family (SREB) with GPR27 and GPR173. These three receptors are “orphan receptors”; however, their endogenous ligands have not been identified. SREB has garnered the interest of many scientists because it is expressed in the central nervous system and is evolutionarily conserved. In particular, brain mass is reported to be increased and learning and memory are improved in GPR85 knockout mice (Matsumoto et al., 2008). In this study, we characterized newly synthesized compounds using a GPR85-Gsα fusion protein and the [^35^S]GTPγS binding assay and identified novel GPR85 inverse-agonists with IC_50_ values of approximately 1 µM. To analyze the neurochemical character of the compounds and investigate the physiological significance of GPR85, we used cerebellar Purkinje cells expressing GPR85 and an electrophysiological technique. Based on the results, the inverse-agonist compound for GPR85 modulated potassium channel opening. Together with the results of previous gene analysis of GPR85, we expect that the development of the GPR85 ligand will provide new insights into a few types of neurological disorders.

## 1. Introduction

The development of new synthetic small molecule ligands is indispensable not only for the development of new drugs, but also for the discovery of new biology and the elucidation of less understood biological phenomena. Human genome analysis has revealed many genes and proteins whose biological functions are not understood. Genetically-modified mice have been mainly used in physiological analysis of these putative gene products. However, for humans, it is natural that genetic modifications are not accepted. Thus, to apply the functional information obtained by genetically modified mice to humans, a pharmacological method using synthetic small-molecule compounds is necessary. G protein-coupled receptors (GPCRs) are transmembrane receptors and are the targets of many clinical drugs. Many synthetic ligands have been developed for GPCRs and have been applied to the functional analysis of target GPCRs and clinical studies. Following our genome analysis of GPCRs (Takeda, Kadowaki, Haga, Takaesu, & Mitaku, 2002), we reported several novel GPCRs ligands, including orphan GPCRs, of which the endogenous ligands and physiological functions remain unknown (Enomoto et al., 2017; Nikaido et al., 2015; Takeda et al., 2003; Uno et al., 2012; Yanai et al., 2016).

GPR85 is mainly expressed in the brain and is an orphan GPCR. Interestingly, GPR85 is the most evolutionarily conserved GPCR and has an identical amino acid sequence in rodents and humans (Hellebrand, Schaller, & Wittenberger, 2000). It was also reported that neurons of the hippocampal dentate gyrus highly expressed GPR85 mRNA in rodents and humans (Matsumoto et al., 2005). One of the key findings of GPR85 studies in genetically modified mice is the involvement of GPR85 in determining brain size. Brain mass was reduced in GPR85 overexpressing transgenic mice and increased in GPR85 knockout mice (Matsumoto et al., 2008). GPR85 overexpression inhibits mouse hippocampal neurogenesis (Chen et al., 2012). In addition, GPR85 knockout mice showed better performance in remembering the conditioning stimulus than their wild-type littermates (Matsumoto et al., 2008). These results indicate that GPR85 is one of the factors influencing the regulation of synaptic plasticity, including learning and memory, and modulating diverse behaviors. For humans, genetic studies have revealed that the GPR85 gene is potentially related to schizophrenia, as single nucleotide polymorphisms (SNPs) in the GPR85 gene are associated with the risk of schizophrenia (Matsumoto et al., 2008). Magnetic resonance imaging (MRI) measurements for detecting volumetric changes in the brain revealed an association between the SNPs and reduced hippocampal volume in patients with schizophrenia, but not in normal volunteers (Radulescu et al., 2013). In addition, possible relationships have been discussed between GPR85 and autism (Trikalinos et al., 2006) (Voineagu et al., 2011), attention deficit hyperactivity disorder (Anney et al., 2008), Tourette’s syndrome, and intellectual disability (Patel et al., 2011). Thus, SNPs in GPR85 might contribute to several neurodevelopmental psychiatric disorders, as well as schizophrenia. Considering these genetic analyses of GPR85, we propose that GPR85 antagonists can be expected to improve a few types of neurological diseases. Owing to their sequence homology and high expression in the brain, GPR27, GPR85, and GPR173 constitute a particular subfamily of GPCRs, namely Super Conserved Receptor Expressed in Brain (SREB). Previously, we reported the identification of several inverse-agonists for SREB using their constitutive activities. The identified ligand compounds showed non-specific activity for three receptors in the SREB subfamily, with individual maximum inhibitory concentrations (IC_50_s) of over 30 µM (Yanai et al., 2016). Since small molecule ligands for SREB have not been previously reported, identification of these compounds provides important insights into the development of SREB ligands. However, their low potency and low water solubility made them difficult to apply to cell experiments to elucidate the physiological function of SREB. It was obvious that the development of new ligands with higher potency was necessary. In this paper, we report the development of new GPR85 ligands that showed inverse-agonist activity at concentrations lower than those of previously reported compounds. We synthesized over 40 chemical compounds to develop compounds with high potency and characterized them using a GPR85-Gsα fusion protein and a [^35^S]GTPγS binding assay (Takeda et al., 2004). The increased [^35^S]GTPγS uptake into Gsα, which was enhanced by the constitutive activity of GPR85 without the addition of an authentic agonist, was reduced by our newly synthesized compounds, with an IC_50_ value of approximately 1 µM. Furthermore, using an electrophysiological technique, we confirmed the activity of the newly identified compound on cerebellar Purkinje cells (PCs), in which we previously reported the expression of GPR85 (Yanai et al., 2016). The results showed that the inverse-agonist compound for GPR85 affected the opening performance of potassium channels in PCs. These results suggest that GPR85 could be a potential target for new therapeutic agents against neurological disorders.

## 2. Materials and Methods

### 2.1. Molecular design and chemical synthesis of the ligand candidate compounds

Based on the chemical skeleton of NPD6061 and NPD6145, which were the SREB inverse-agonists previously reported (Yanai et al., 2016), we focused on the two aromatic rings and the chemical bond connecting them as pharmacophore structures (Fig. 1). The basic molecular skeleton consisted of one benzene ring with a hydroxyl or methoxy group at the ortho position, the other benzene ring with a hydroxyl group at the para position, and a linker with three atoms. Molecules with such a molecular skeleton were designed and synthesized using amide or ester bonds. Amide and ester compounds are suitable for obtaining information on structure-activity relationships in the early stages of ligand development; this is because they are easy to synthesize and can be used to create various compounds in a short period by combining a carboxylate portion with an alcohol portion.

**Figure 1.**
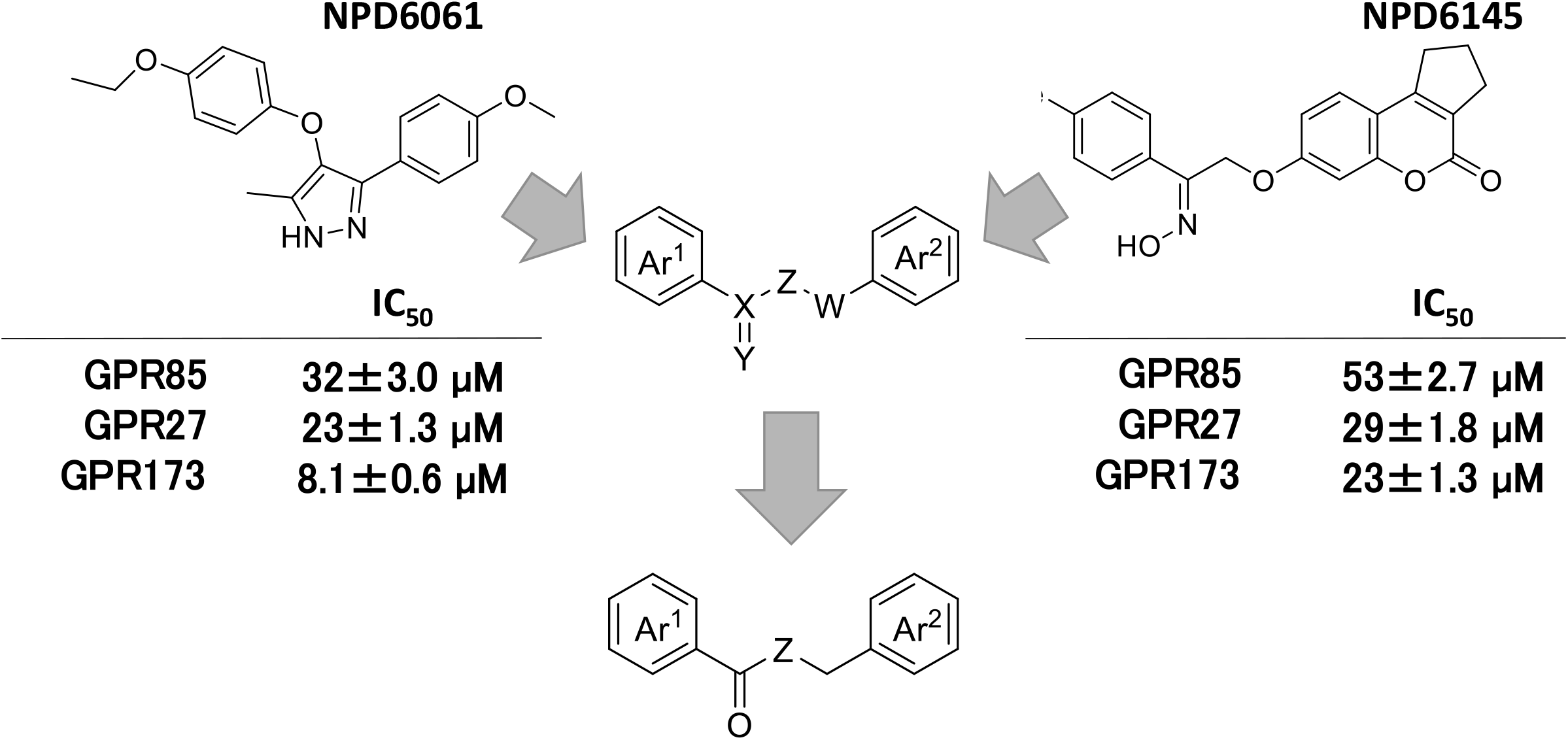
Molecular design of novel ligands for GPR85 based on previously-reported inverse-agonists with with their IC_50_s (Yanai et al., 2016) and focused pharmacophore structures. We noticed that NPD6061 and NPD6145 consist of a molecular skeleton in which two benzene rings are linked by a linker with three atoms.

Amide derivatives **1a, 1b**, and **1c** were synthesized by condensation reactions using 2-hydroxy-4-methoxybenzoyl chloride and the corresponding amines in the presence of ^*i*^Pr_2_NEt as a base (Fig. 2). Most ester derivatives were also prepared from the corresponding benzoyl chlorides and alcohols in a similar manner to the synthesis of amide derivatives. Ester derivatives **2a**–**2h** and **3a**–**3k** (Fig. 3) were obtained by condensation of the corresponding benzoic acids and alcohols using 1-ethyl-3-(3-dimethylaminopropyl)carbodiimide hydrochloride (EDC · HCl) as a condensation reagent. All compounds were purified by silica gel column chromatography using hexane and ethyl acetate as eluents. The structure of the purified product was confirmed by ^1^H-NMR, ^13^C-NMR, and TOF mass spectrometry.

**Figure 2.**
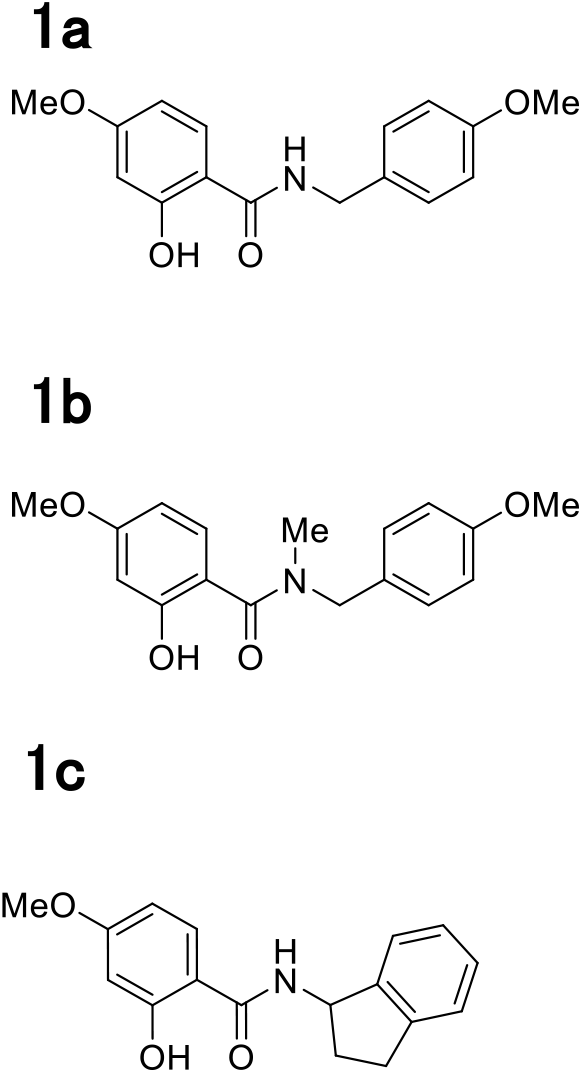
Newly-synthesized amide derivatives for the GPR85 ligand candidates.

**Figure 3.**
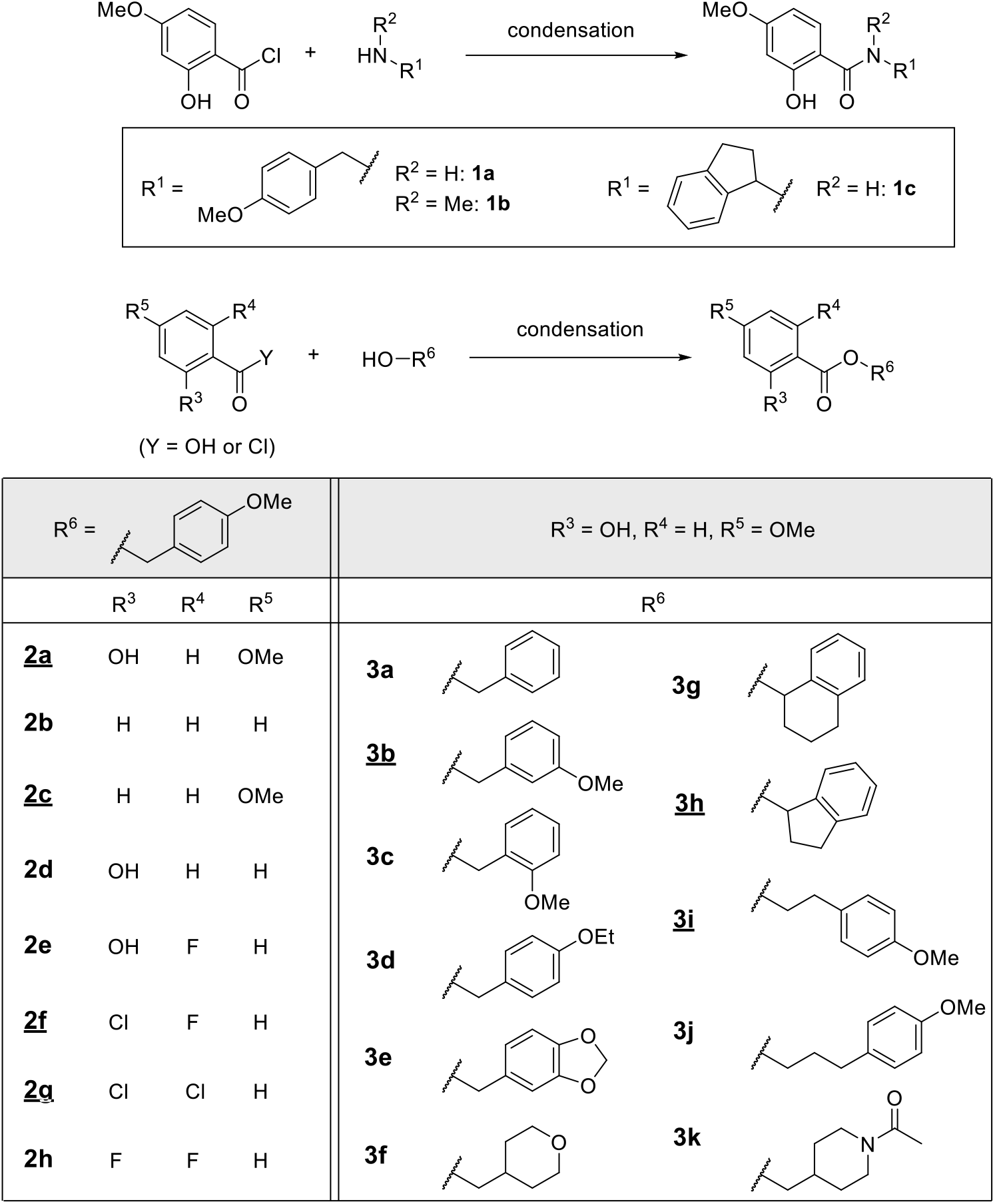
Newly-synthesized ester derivatives for the GPR85 ligand candidates. The underlined compounds showed significant inverse-agonist activity for a GPR85-Gα fusion protein at 10 µM.

### 2.2 *In vitro* [^35^S]GTPγS binding assay

Characterization of ligand activity based on SREB-induced [^35^S]GTPγS uptake into Gα was performed using an SREB-Gsα fusion protein as previously described (Takeda et al., 2004). Briefly, the SREB-Gsα fusion gene was constructed using a two-step PCR. The resulting fusion gene was used to prepare the transgene virus to infect Sf9 cells using the Bac-to-Bac system. After infecting Sf9 cells with the recombinant virus for 96 h, Sf9 cells expressing an SREB-Gsα fusion protein on the cell membranes were collected. Thereafter, a cell membrane suspension was prepared and used for further activity measurement. For the ligand assay, a cell membrane suspension (20 µg of total protein) and a ligand compound were added to a reagent solution (20 mM HEPES-KOH [pH 8.0], 1 mM EDTA, 160 mM NaCl, 1 mM dithiothreitol, 100 pM [^35^S]GTPγS (PerkinElmer, MA, USA), 1 µM Na-GDP, and 10 mM MgCl_2_, with a final volume of 100 µL). The reaction mixture was incubated at 30 °C for 30 min, and then filtered through a GF/B glass fiber filter (Whatman, Kent, UK) to trap the cell membrane to which [^35^S]GTPγS was bound. The amount of radiation remaining on the glass fiber filter was measured using a liquid scintillation counter (Tri-Carb 2910TR, PerkinElmer). For the entire *in vitro* assay, the synthesized ligands were added to the reaction solution at a final concentration of 10% DMSO. As this [^35^S]GTPγS binding assay is applicable to a synthetic chemical dissolved in DMSO, it is suitable for measuring the activity of a lead compound with low water solubility in the early stages of drug discovery. The resultant data points were fitted using KaleidaGraph (HULINKS, Japan) to plotted a curve to estimate the IC_50_ (Yanai et al., 2016).

### 3.3 Electrophysiological experiments

Electrophysiological experiments were performed as previously described (Watanave et al., 2019) (Watanave et al., 2018). Parasagittal cerebellar slices (250 μm in thickness) were prepared from 7-week-old C57BL6/J mice. Slices were perfused in extracellular solution (125 mM NaCl, 2.5 mM KCl, 1.25 mM NaH_2_PO_4_, 26 mM Na_2_HCO_3_, 2 mM CaCl_2_, 1 mM MgCl_2_, 10 mM glucose, bubbled continuously with a mixture of 95% O_2_ and 5% CO_2_) at room temperature during the recordings. The resistance of the patch pipette was 2 – 5 MΩ when filled with intracellular solution (122.5 mM Cs-gluconate, 17.5 mM CsCl, 8 mM NaCl, 2 mM Mg-ATP, 0.3 mM Na-GTP, 10 mM HEPES-CsOH, 0.2 mM EGTA).

PCs were clamped at ‒70 mV to record the excitatory postsynaptic currents (EPSCs) of parallel fiber (PF). Stimulation pipettes were filled with the extracellular solution and placed in the molecular layer to activate the PFs. PF EPSCs were monitored every 10 s. We confirmed the stability of the PF EPSCs for at least 10 min before drug application. The series resistance was continuously monitored every 10 s by applying small hyperpolarizing pulses. Data were discarded when the resistance values changed by >20% of the basal value during the course of the experiment. Selective stimulation of PFs was confirmed by paired-pulse facilitation (PPF) of the EPSC amplitudes with a 50-ms inter-stimulus interval (ISI). The PPF ratios of each trace were calculated by dividing the 2nd EPSC amplitudes by the 1st EPSC amplitudes. Amplitudes of PF EPSCs and PPF ratios were averaged every minute (six traces) and normalized by the average value of the responses over 10 min immediately before the bath application of **3i** (0.1 mM). The extracellular solution for EPSC recording contained 0.1 mM picrotoxin.

Passive currents were measured from PCs in the presence or absence of a ligand compound (0.1 mM **3i** or 0.1 mM **1c**). The intracellular solution used to record passive currents contained 122.5 mM K-gluconate, 17.5 mM CsCl, 8 mM NaCl, 10 mM HEPES, 0.2 mM EGTA, 2 mM Mg-ATP, 0.3 mM Na-GTP (pH 7.2, 290-300 Osm). PCs were first held at -70 mV, and then currents were evoked by moving the holding potentials to various potentials for 1 s. The currents obtained from the last 50 ms were estimated as the passive currents. One-way ANOVA followed by Bonferroni post-hoc comparisons were used to analyze the data. Differences were considered significant at P < 0.05.

## 3. Results

As we previously reported that both NPD6061 and NPD6145 have two aromatic rings as pharmacophores, we derived a strategy to connect aromatic rings with different substituents to develop new ligand compounds (Fig. 1). Benzyl benzoate and *N*-benzylbenzamide were assumed to be the basic skeleton, and more than 40 benzoate and benzamide derivatives, including benzyl benzoates, were synthesized by the condensation of substituted benzoic acids and various alcohols or amines. No further studies were performed on the compounds that showed no activity when the [^35^S]GTPγS binding assay was carried out at a concentration of 100 µM. At this stage, the chemicals with an amide bond (**1a, 1b, 1c**) did not exhibit ligand activity and the ligand candidates were composed of an ester bond (Fig. 2). To investigate the compounds that showed activity at 10 µM, the compounds shown in Figure 3 were systematically synthesized, their [^35^S]GTPγS binding activities were measured, and their structure-activity relationships were examined. We observed significant inverse-agonist activity with **2a, 2c, 2f, 2g, 3b, 3h**, and **3i** for a GPR85-Gα fusion protein at 10 µM (Fig. 3, Fig. S1). Fortunately, the newly synthesized **2g, 3h**, and **3i** showed inverse-agonist activities for GPR85 in the [^35^S]GTPγS binding assay at lower concentrations than those previously reported, NPD6061 and NPD6145 (Fig. 4). The IC_50_ values were 1.7 ± 0.2 µM, 1.3 ± 0.4 µM, and 0.52 ± 0.3 µM, for **2g, 3h**, and **3i**, respectively (Table I). As inverse-agonist activities were also observed for GPR27 and GPR173 in the [^35^S]GTPγS binding assay, we concluded that they were non-selective ligands for the SREB subfamily (Fig. 5). The IC_50_ values for GPR27 were 2.3 ± 0.5 µM, 1.5 ± 0.6 µM, and 0.67 ± 0.3 µM, while those for GPR173 were 2.6 ± 0.9 µM, 1.1 ± 0.7 µM, and 2.6 ± 0.9 µM, for **2g, 3h**, and **3i**, respectively (Table I). Essentially, the three compounds showed almost the same potency for the three SREB receptors. It is a great advantage for the [^35^S]GTPγS binding assay that the ligand activity is measured even with 10% DMSO. It has been reported that the ligand affinity obtained using the isotope-labeled compounds and the potency obtained using the GPCR-Ga fusion proteins gave a good correlation (Takeda et al., 2004; Zhang, Okamura, Guo, Niwa, & Haga, 2004). In this result as well, it was reasonable to suppose that the potency correlated with the binding affinity, and that **2g, 3h**, and **3i** with lower IC50s were expected to have higher affinities to GPR85.

**Figure 4.**
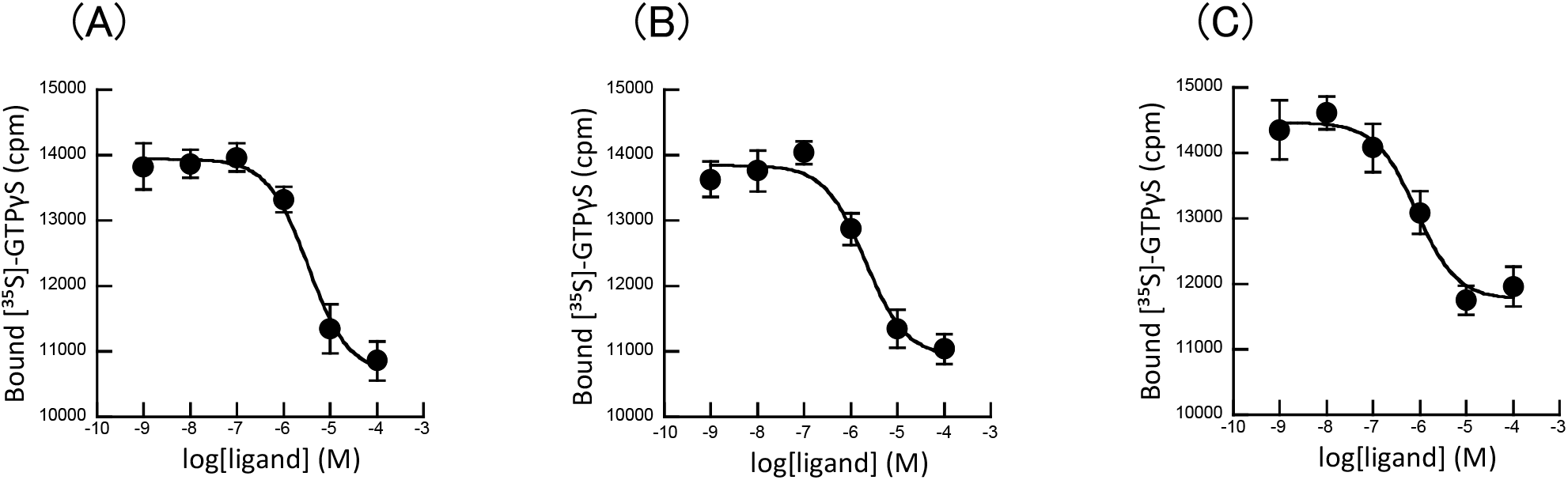
Characterization ligand compounds in the [^35^S]GTPγS binding assay for GPR85, (A); **2g**, (B); **3h**, (C); **3i**. Each data point is the mean ± SEM of 3 measurements. Their IC_50_ values are summarized in Table I.

**Table I.**
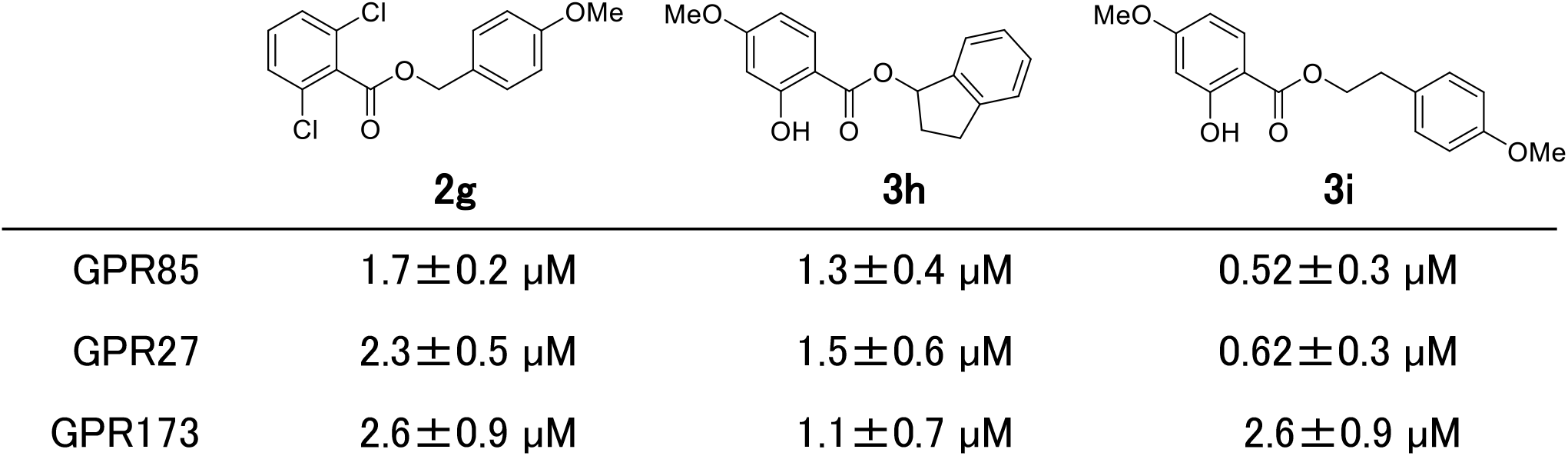
Summary of the potency of identified inverse agonists for SREB-Gsα fusion proteins measured using the [^35^S]GTPγS-binding assay. Each value represents the mean ± SEM of 3 independent experiments.

|  |  |  |  |
| --- | --- | --- | --- |
| GPR85 | $1.7 \pm 0.2 \text{ } \mu\text{M}$ | $1.3 \pm 0.4 \text{ } \mu\text{M}$ | $0.52 \pm 0.3 \text{ } \mu\text{M}$ |
| GPR27 | $2.3 \pm 0.5 \text{ } \mu\text{M}$ | $1.5 \pm 0.6 \text{ } \mu\text{M}$ | $0.62 \pm 0.3 \text{ } \mu\text{M}$ |
| GPR173 | $2.6 \pm 0.9 \text{ } \mu\text{M}$ | $1.1 \pm 0.7 \text{ } \mu\text{M}$ | $2.6 \pm 0.9 \text{ } \mu\text{M}$ |

**Figure 5.**
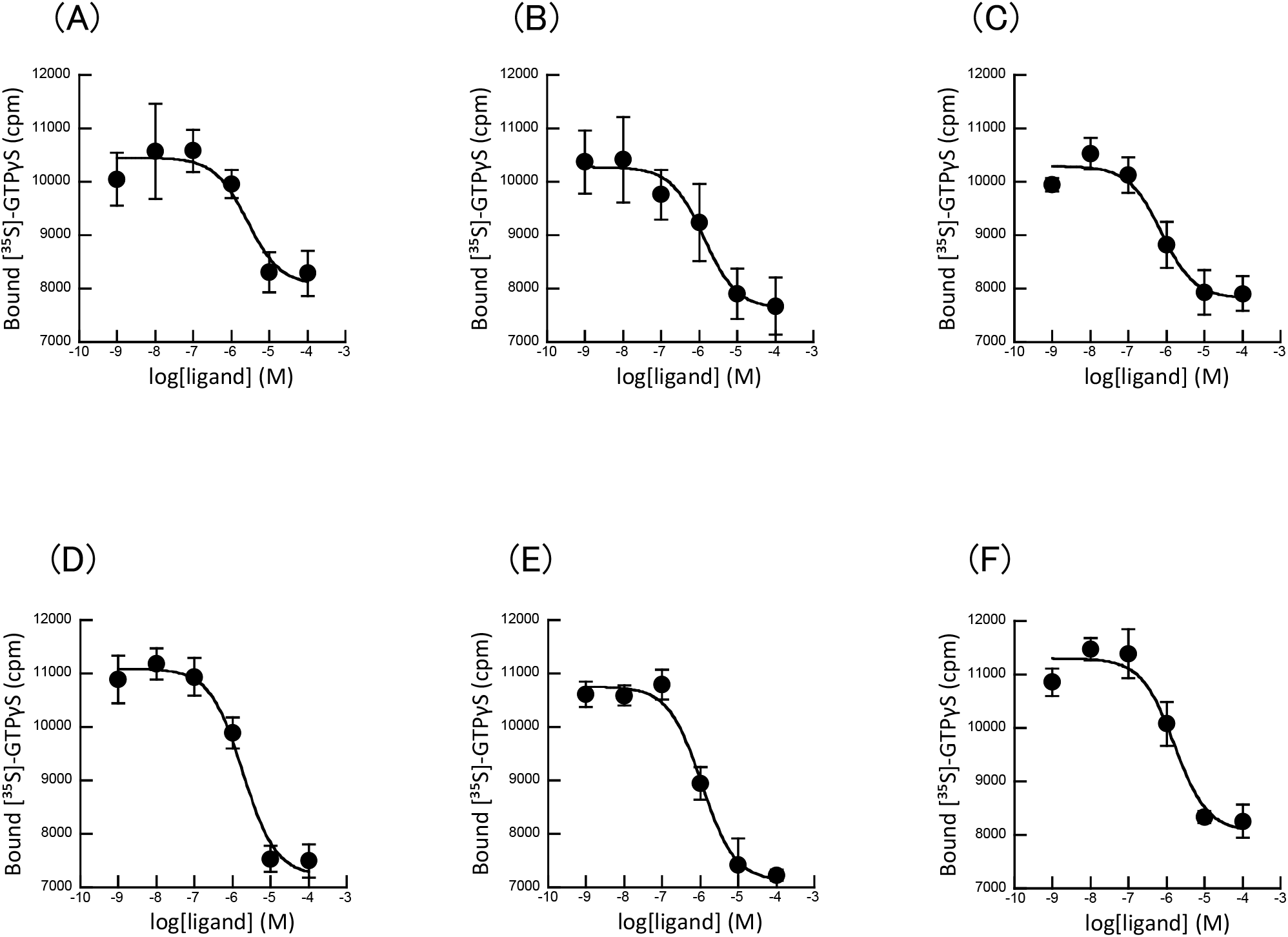
Characterization ligand compounds in the [^35^S]GTPγS binding assay for GPR27, (A); **2g**, (B); **3h**, (C); **3i**, for GPR173, (D); **2g**, (E); **3h**, (F); **3i**. Each data point is the mean ± SEM of 3 measurements. Their IC_50_ values are summarized in Table I.

Electrophysiological experiments using nerve cells of brain slices were extremely sensitive to organic solvents and could be studied using water-soluble compounds. Owing to their water solubility, we examined **3i** as an active ligand and **1c** as a negative control compound. As **2g** and **3h** precipitated within a few hours after being dissolved in the experimental buffers, their water solubility was poor. As a result, they could not be used in the electrophysiological experiments. In contrast, **1c**, a compound with an amide bond, maintained water solubility. Only slight differences were found in the chemical structure, but there were serious alterations and improvements not only in biological activity, but also in the physical properties, such as water solubility, among **1c, 2g, 3i**, and **3h**. For the electrophysiological evaluation of **3i**, we first examined EPSCs elicited at PCs by electrical stimulation of PFs (PF-EPSCs) (Fig. 6A). Bath-application of **3i** or **1c** did not influence the PF-EPSC amplitude (Fig. 6B and 6C), suggesting that **3i** and **1c** had no acute neurotoxicity and did not affect basal synaptic transmission between PF and PC. We also measured the PPF ratios to determine the effect of **3i** or **1c** on the presynaptic release probability from PFs. We found no significant difference in PPF ratios with or without **1c** or **3i** (Fig. 6D and 6E). These results indicated that GPR85 and its inverse agonist did not influence basal synaptic transmission or short-term synaptic plasticity. Next, we measured the passive currents from the PCs in the presence or absence of **3i** or **1c**. We found that the bath-application of **3i**, but not **1c**, significantly increased the passive current of PCs only when the membrane voltage of PCs became approximately -100 mV or less (Fig. 7). As the passive currents were enhanced only at membrane voltages below -100 mV, **3i** may have augmented the permeability of potassium through the PC membrane (likely enhancing the function of potassium channels), because the equilibrium potential of potassium ions (but not other ions) was very close to -100 mV. Our results of the [^35^S]GTPγS binding assay showed that **3i** worked for all SREB members, namely GPR27, GPR85, and GPR173, and was a non-selective inverse-agonist. However, as we have already reported that only GPR85 in SREB is expressed in PCs, it is reasonable to consider that the effect of **3i** on potassium channels is mediated by GPR85. Collectively, we concluded that the inverse-agonist, **3i**, reduced the constitutive activity of GPR85, leading to functional enhancement of the potassium channels. At the same time, this result indicated that the constitutive activity of GPR85 had an inhibitory effect on the potassium channels in PCs.

**Figure 6.**
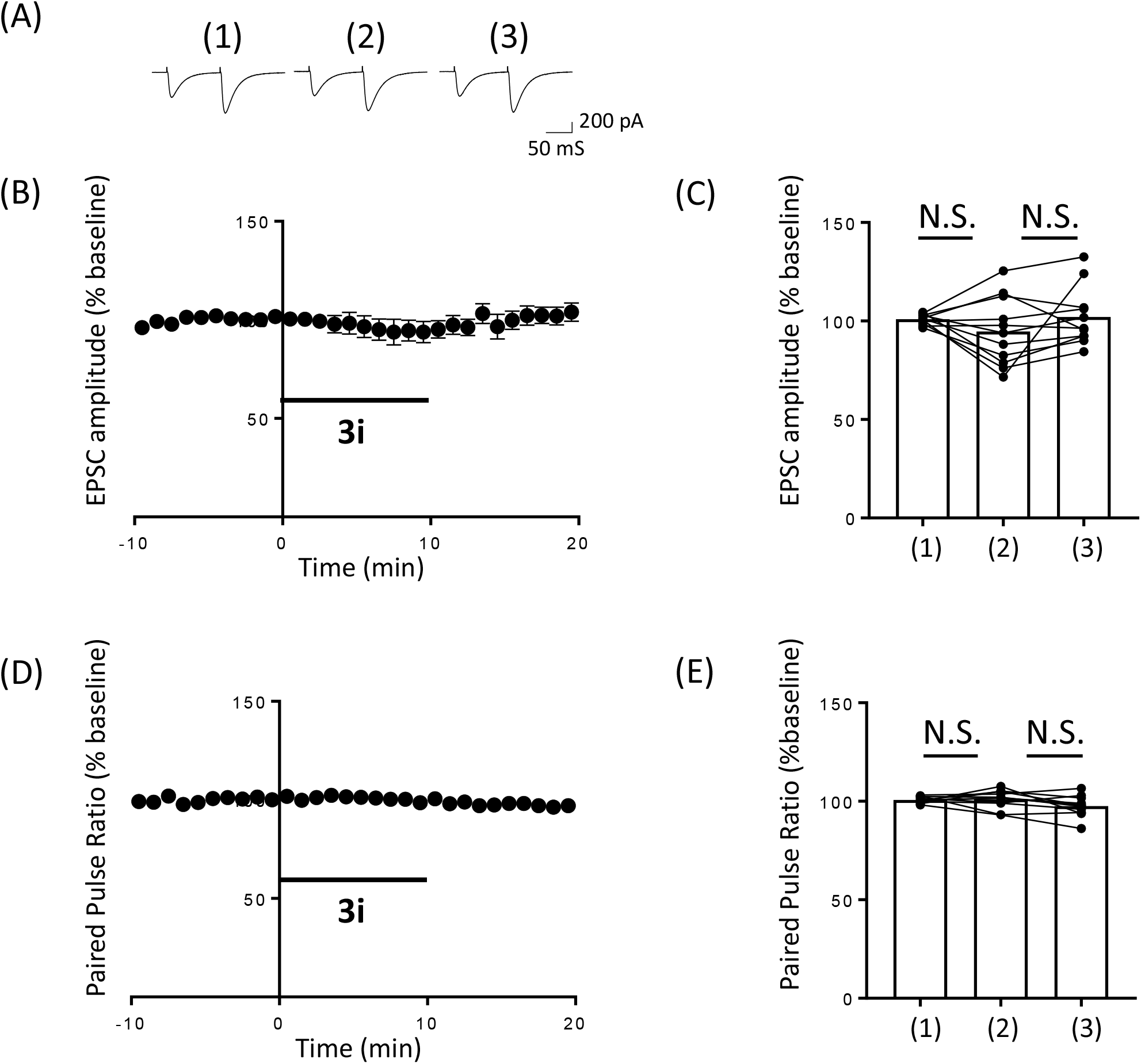
Observations of PPF and evaluation of GPR85 inverse-agonist effects for PF and PCs. The average of the EPSC amplitudes did not show significant difference between the presence and absence of **3i** or **1c**. (A); Representative observation of the PPF measurements. (1); baseline period, (2); in the presence of 0.1 mM **3i**, (3); washout period. (B); Representative results of the EPSC amplitudes. (C); Comparison of the EPSC amplitudes with or without **3i**. (1); baseline period, (2); in the presence of 0.1 mM **3i**, (3); washout period. (D); Representative results of the PPF ratios. (E); Comparison of the PPF ratios with or without **3i**. (1); baseline period, (2); in the presence of 0.1 mM **3i**, (3); washout period.

**Figure 7.**
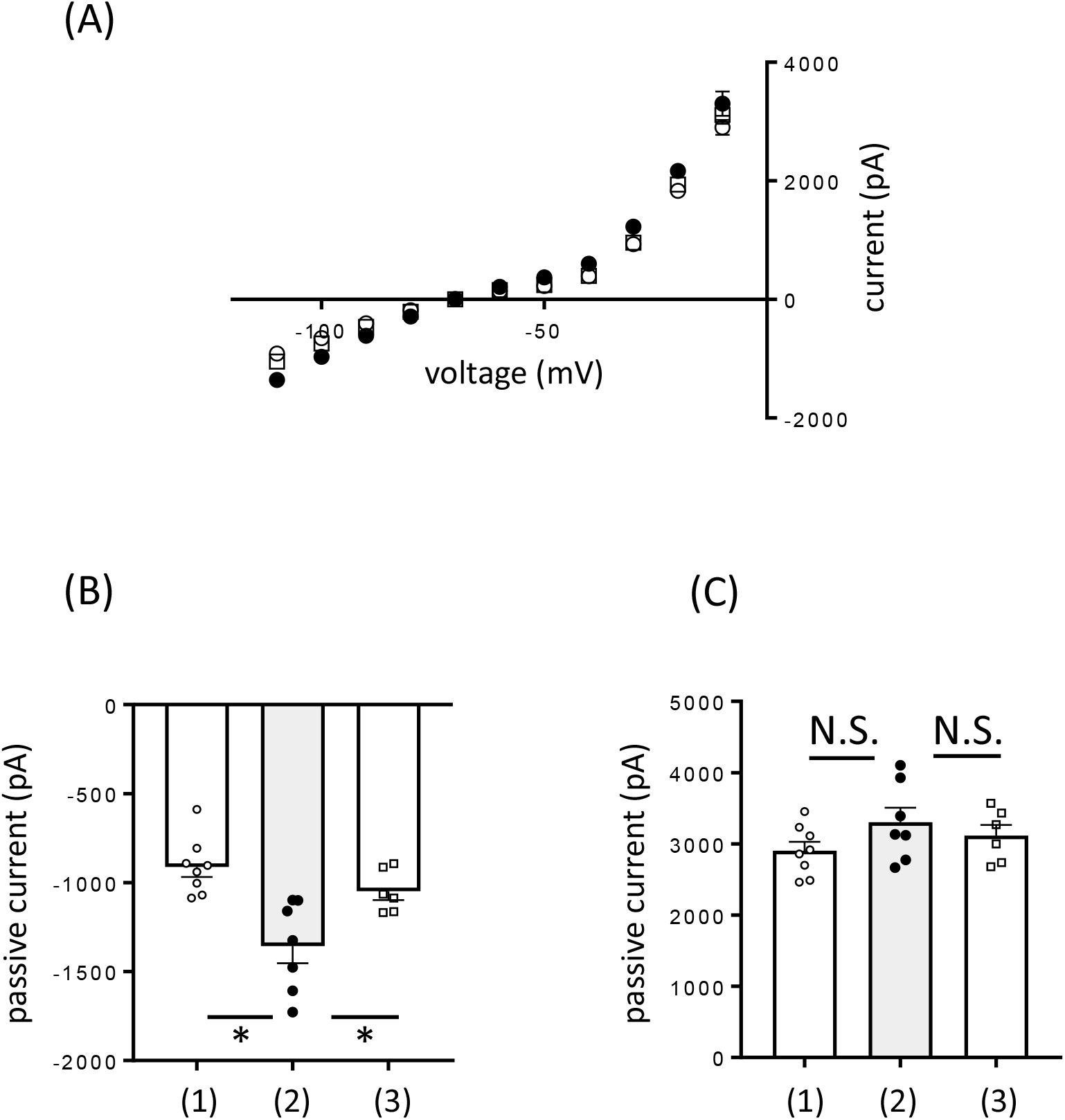
Measurements of the passive current and evaluation of GPR85 inverse-agonist effects for PCs. A significant increase in the measured currents was observed only under the hyperpolarized conditions in the presence of **3i**. (A); Comparison of the *I*–*V* plots with or without **3i** or **1c**. Open circles; normal Ringer solution, closed circles; in the presence of 0.1 mM **3i**, open squares open; in the presence of 0.1 mM **1c**. (B); average of the measured currents clamping at -100 mV, (1); normal Ringer solution, (2); in the presence of **3i**, (3); in the presence of **1c**. Significant enhancement (*P* < 0.05) of the measured current was found for **3i**, but not for **1c**. (C); average of the measured currents clamping at -10 mV, (1); normal Ringer solution, (2); in the presence of **3i**, (3); in the presence of **1c**.

## 4. Discussion

Structure-activity relationship-based organic synthesis and derivative development approaches have been the most reliable and effective method for ligand development, even today when computational ligand development is possible. The fact that **1a, 1b**, and **1c** did not show any activity for GPR85 indicated an amide bond was not acceptable for ligand activity. The reason for the lack of support for an amide bond is currently unclear; however, future studies will demonstrate in detail how ligand compounds exhibit their activity and how they interact with the receptors at atomic resolution.

In our study, benzyl benzoate was found to be a very effective and useful basic skeleton (Fig. 3). Comparing the active **2a** and **2c** having IC_50_s over 5 µM (Fig. S1) with inactive **2b** and **2d**, the methoxy group at the 4-position (R^5^) was supposed to be important for their binding to GPR85. We also investigated the effect of the substituent at the 2- and 6-positions (R^3^ and R^4^) of the benzoic acid portion. With introducing chlorine atom into R^3^, **2f** and **2g** became active. The results that **2f** showed less activity than **2g** indicated that fluorine with a small atomic radius was not large enough. Together with these facts, it was concluded that the presence of a sufficiently large atom at the position of R^3^ and R^4^ had a great effect on activity. On the alcohol portion, the activity of the benzyl alcohol ester (**3a**) was attenuated, confirming that the methoxy group on the aromatic ring was important. Moreover, the spatial arrangement of the methoxy group gave a great influence on the activity. The activity showed a significant difference between the *meta-*position (**3b**) and the *ortho*-position (**3c**) at the aromatic ring. The change in the length of the linker (from **3i** to **3j**) also induced change of the activity. In addition, the activity was lost when the methoxy group at the *para*-position was converted to lager atomic group (**3d** and **3e**). All of these results indicated that the ligand-binding pocket of GPR85 where the alcohol portion bound was very constrained. The fact that **3h** showed excellent activity is reasonable with this consideration because **3h** has a fixed conformation and does not change the conformation in the pocket. Probably because of the limited volume of the binding pocket, **3g** was not accepted in contrast with **3h** and **3k**. In addition, the involvement of a π-electron-related interaction between a ligand molecule and a GPR85 side chain has been suggested, as a non-aromatic ring was not accepted for the alcohol portion (**3f**). Since a few slight differences in the substituents on the benzene ring caused a clear difference in their activities, for example, **2f, 2g, 2h** or **3g, 3h, 3i, 3j**, we concluded that the entire molecular structure of the small ligand identified herein, **2g, 3h**, and **3i** interacted closely with GPR85 and were positioned and inserted into the ligand pocket of GPR85. Preliminary docking model simulations supported this conclusion (Fig. S2). As all three compounds had no selectivity and showed almost the same activity for GPR27, GPR85, and GPR173, the structures of the ligand pocket of the three receptors were suggested to be very similar. We finally improved the IC_50_ to approximately 60-fold that of previously reported compounds. Further, development of the SREB ligand is steadily being achieved.

We were able to apply **3i** to electrophysiological experiments because of its appropriate potency and water solubility. As **3i** did not affect the ratio of the PPF at the PF to PC synapses, we surmised that GPR85 was not involved in glutamate release from the PF terminals (Fig. 6). Although improved memory was observed in GPR85 knockout mice (Matsumoto et al., 2008), the analysis with the GPR85 inverse-agonist, *i*.*e*. **3i**, did not provide evidence regarding the contribution of GPR85 to short-term synaptic plasticity. In contrast, **3i** was found to promote potassium channel function in electrophysiological experiments using mouse cerebellar slices (Fig. 7). Such finding indicates that GPR85 acts on potassium channels and highlights our success at developing a reagent, which is applicable for neuroscience research, to overcome water solubility and toxicity issues. Moreover, to our knowledge, this is the first report to outline the molecular mechanism of GPR85 in physiological action. The G protein-gated inward-rectifying potassium channels are described to be potential regulators of neuronal excitability and are regulated by multiple GPCRs, typically in opioid receptors (Nagi & Pineyro, 2014). These channels are usually activated by the Gβγ complex. However, in practice, it has also been reported that there are various mechanisms for the interaction between potassium channels and GPCRs. Some potassium channels have been shown to interact directly with Gβγ and Gα (Oldham & Hamm, 2006) (Johnston & Siderovski, 2007). A similar current increase in the hyperpolarized state by an inverse-agonist function was previously reported for melanocortin 4 receptor (MC4R) (Ghamari-Langroudi et al., 2015). Agouti-related protein (AgRP) is an endogenous inverse agonist that blocks the constitutive activity of MC4R. The regulation of hypothalamic neuron firing activity by AgRP has been reported to be mediated by ligand-induced coupling of MC4R to regulate inward-rectifying potassium channels. Inhibition of MC4R by the endogenous inverse agonist, AgRP, caused neuronal channel opening and hyperpolarization. In contrast, activation of MC4R resulted in powerful neuronal depolarization, which was found to be mediated by potassium channel closure. As our results revealed that GPR85 could modulate potassium channels, we expect that potassium channel regulation by GPCRs would be analyzed in detail by developing various SREB ligands as well as MC4R and AgRP.

In the near future, we plan to carry out *in vivo* experiments using animal models. However, we expect that our ester compounds will be easily hydrolyzed inside the animal body and will maintain their activity. Therefore, it is necessary to develop other compounds with appropriate *in vivo* stability. We also aim to design a ligand that permeates the blood-brain barrier but has sufficient water solubility. We presume that compounds with an ether structure instead of ester would be one of the candidates with high activity and appropriate pharmacokinetics. Neutral antagonists prevent agonists from binding, but they do not shift the equilibrium between active and inactive receptors and do not alter the basal level of signal transduction. However, inverse agonists preferentially bind to the inactive state of the receptor, reducing the proportion of active receptor and suppressing basal signaling including constitutive activity. Inhibition of nerve cell activation by GPR85 inverse agonists will lead in a pharmacologically similar situation to GPR85 deficiency in knockout mice. It would be interesting to determine whether the phenomena observed in GPR85 knockout mice, for example, improved learning and memory, can be reproduced by halting the GPR85 functions with our inverse agonists.

## Acknowledgments

This research was partially supported by the Platform Project for Supporting Drug Discovery and Life Science Research (Basis for Supporting Innovative Drug Discovery and Life Science Research (BINDS)) from AMED under Grant Number JP21am0101099 (support number 0950). This work was also supported by a grant from the Gunma University Medical Innovation Project to H.H. and S.T. The cell membranes expressing an SREB-Gsα fusion protein were collected using an ultracentrifuge at the Center for Instrumental Analysis of Gunma University.

